# Ciliopathy-associated missense mutations in IFT140 are hypomorphic and have edgetic effects on protein interaction networks

**DOI:** 10.1101/2023.01.18.523235

**Authors:** Tobias Leonhard, Gaurav D. Diwan, Franziska Klose, Isabel F. Stehle, Katrin Junger, Marian Seda, Sylvia Bolz, Franziska Woerz, Robert B. Russell, Karsten Boldt, Dagan Jenkins, Marius Ueffing, Tina Beyer

**Author notes:** Authors for correspondence: Tina Beyer [ ]; Dagan Jenkins.

## Abstract

The mechanisms underlying recessive Mendelian diseases and the interplay between genotype and phenotype still need to be better understood. It is therefore necessary to characterise the functional effects of missense mutations at the protein level. Here we focus on missense mutations in the intraflagellar transport protein IFT140, which forms part of the IFT complex A (IFT-A), a crucial component of the ciliary machinery. Mutations in IFT140 can cause a vast spectrum of diseases belonging to the group of ciliopathies, reaching from isolated retinal dystrophy to severe skeletal abnormalities and multi-organ diseases such as Mainzer-Saldino and Jeune syndrome. We hypothesise that missense mutations in IFT140 are hypomorphic leading to quantitative effects on a subset of protein-protein interactions. This may affect complex stability as well as perturbations of protein interaction networks. In this work we assessed how 24 missense mutations in IFT140 affect interactions with other IFT and effector proteins using affinity purification coupled to mass spectrometry. Our data reveals that several mutations in IFT140 are hypomorphic and disrupt the stability of the IFT-A complex to varying degrees in a quantitative way. Allelic combination and the degree of IFT-A complex disruption in analysed missense mutations correlates with the severity of the observed phenotype in a subset of patients. In addition, we show that a distinct subset of mutations in IFT140 shows edgetic effects by disrupting specific PPIs rather than causing a total loss of IFT-A binding. This is the case e.g. with the disease-associated protein TULP3 which is involved in cilia-dependent sonic hedgehog signalling.

## Introduction

Understanding the complex interplay between genotype and phenotype in inherited disease and the underlying pathogenic mechanisms remains a major challenge. Potential mechanisms include e.g. abundance of functional protein, aberrant PTMs, disruption of protein complexes as well as the alteration of specific PPIs. The perturbation of a specific PPI is referred to as edgetic effect, with genetic defects in recessive-mendelian disease leading to the loss of a specific connection (edge) in an interaction network rather than the complete removal of one component, or node, within this network (Del-Toro et al. 2019). While disease mechanisms for specific mutations have been identified in various studies (Badonyi and Marsh 2022; N. Sahni, M. Taipale 2017), analysis of large numbers of disease-associated mutations for one specific gene are rare. In this study we aimed to better understand the genotype-phenotype correlation and the corresponding molecular mechanisms by analysing the impact of 24 disease-associated missense mutations in the ciliary protein IFT140. By stable overexpression of IFT140 wildtype and mutant constructs in HEK293 cells followed by affinity purification and MS/MS analysis we assessed the changes in protein interaction networks induced by these mutations.

Cilia are antenna-like organelles protruding from the surface of most polarised eukaryotic cells, which are highly conserved throughout evolution (Avasthi and Marshall 2012). They play a crucial role in a broad variety of vital processes, including neurosensory functions such as hearing and vision, reproduction, tissue development and homeostasis (Reiter and Leroux 2017). Primary cilia usually serve as sensing organelles and signalling platforms and they are a key component for signalling pathways such as sonic hedgehog (Anvarian et al. 2019; Liem et al. 2012).

Transport of cargo such as proteins within the cilium is referred to as intraflagellar transport (IFT). It involves motor proteins as well as protein complexes, which enable specific cargo selection through several protein-protein interaction (PPI) motifs (Boldt et al. 2011). IFT is crucial for the proper assembly, maintenance and function of eukaryotic cilia. The high degree of evolutionary conservation of the IFT particles in eukaryotic cells hints towards the critical role of each single component for the proper function of cilia.

Retrograde transport from the ciliary tip to the ciliary base of cargo like GPCRs is achieved by active transport with dynein 2. For mediating contact between motor protein and cargo a protein complex called IFT-A is necessary. IFT-A consists of six subunits and can be further distinguished into two subcomplexes (Taschner, Bhogaraju, and Lorentzen 2012). IFT140, IFT122 and IFT144 form the core complex, which is attached to a peripheral subcomplex consisting of IFT43, IFT121 and IFT139 (Mukhopadhyay et al. 2010; Hesketh et al. 2022). Due to the large size of some of the subunits and the sensitivity of the complex to stoichiometric changes, elucidating the precise role and function of several of these components has proven challenging in the past. IFT-A also has functions in the anterograde transport, e.g. by linking heterotrimeric kinesin to its corresponding IFT particle, which enables its transport from the ciliary base to the ciliary tip (Ou et al. 2007; Nakayama and Katoh 2018; Mukhopadhyay et al. 2010; Nakayama and Katoh 2020).

IFT140 is a core components of IFT-A and plays a crucial role in the proper function of the ciliary machinery (Crouse et al. 2014). The function of IFT140 includes not only retrograde transport of cargo within the cilium but also the ciliary entry of several cargo proteins such as GPCRs (Hirano et al. 2017; Mukhopadhyay et al. 2010). Defects in IFT140 can lead to a vast spectrum of diseases, which are commonly referred to as ciliopathies. These disorders are inherited through a recessive-mendelian mechanism, meaning that both alleles must be affected to cause disease. Biallelic mutations in the IFT140 coding gene can affect different tissues and to varying degree (Perrault et al. 2012a; Khan, Bolz, and Bergmann 2014a; Hull et al. 2016). The milder forms include isolated retinal dystrophies as well as retinitis pigmentosa (RP) and Leber congenital amaurosis (LCA). They all cause vision loss in affected individuals, but severity and age of onset can vary significantly. While LCA usually presents itself at birth or during early infancy the onset of retinitis pigmentosa can be as late as mid-adulthood (Xu et al. 2015a; Hull et al. 2016). The more severe forms cause skeletal ciliopathies like Jeune asphyxiating thoracic dystrophy and Mainzer-Saldino syndrome. One of the main features of this disease are shortened ribs, leading to a narrow thorax. This restricts lung growth and function and frequently causes perinatal death due to asphyxia. More variable features include polydactyly, polycystic kidney disease, retinal dystrophy and liver disease (Perrault et al. 2012a; Oud et al. 2018). Mainzer-Saldino syndrome is defined by phalangeal cone-shaped epiphyses in combination with renal disease and retinal dystrophy. Less common features include cerebellar ataxia and liver fibrosis as well as short stature (Perrault et al. 2012b). Severe defects in IFT140 have been shown to be lethal during embryonic development, further highlighting the crucial role of functional IFT140 (Perrault et al. 2012a). We therefore hypothesize that the missense mutations in IFT140 reported in patients suffering from ciliopathies are hypomorphic and only reduce native protein function to a certain degree, defining type and severity of the disease.

By stable overexpression of IFT140 wildtype and mutant constructs fused to an N-terminal Strep/FLAG-tag we examined the interactome using two approaches: affinity purification followed by LC-MS/MS analysis to determine interactome changes and co-immunoprecipitation to validate changes in specific PPIs. Our results tie several mutations to quantitative changes in specific protein interactions which confirmed the hypothesis of hypomorphic and edgetic effects of IFT140 missense mutations.

## Results

### Identification of the IFT140 wildtype interactome

In a first step we aimed to gain further insight into the PPIs of wildtype IFT140 (IFT140 WT) and to assess the robustness of our approach. Using HEK293 cells stably expressing IFT140 WT or RAF1 as negative control, both N-terminal fused to a Strep/FLAG-tag, a Strep affinity purification followed by LC-MS/MS analysis and labelfree quantification was performed and the interactome of wildtype IFT140 was determined (Figure 1). Using a total of 36 biological replicates from 12 separate experiments we were able to obtain an extremely robust dataset for statistical analysis. The labelfree quantification was done using the MaxQuant software and statistics were calculated using R script. Proteins were classified as interactors if they were enriched with a corrected Student’s t-test p-value <= 0.05 and significance A value <= 0.05 in the IFT140 samples as compared to the RAF1 control. Figure 1 shows a scatter plot representing the peptides retrieved, and proteins that are enriched following IFT140 pulldown are labelled. 15 proteins were found to be enriched in our data (Figure 1, Table 1), including all components of the IFT complex A. This proves the robustness of our approach and its suitability for interactome identification. Besides the IFT-A complex itself, known interactors of IFT-A were enriched, such as the ciliary protein TULP3, a negative regulator of hedghehog signalling. SYNJ2 was also found as enriched, a protein involved in membrane trafficking and signal transduction (Chuang et al. 2012). Another previously reported interactor of IFT140 is NUDC, which is involved in spindle formation during mitosis and in microtubule organization during cytokinesis. Gene enrichment analysis of the generated protein-protein interaction data using STRING (https://string-db.org/) revealed 8 so far unknown potential interactors as enriched in our data (Figure 1). To illuminate potential new roles of IFT140 the function of each so far unknown potential interactor has been searched (Table 2). PLEC has been proposed to be involved in the organization of the cytoskeleton, where it interlinks intermediate filaments with microtubules (Wiche 1998). Apart from its role in the nonsense-mediated decay of mRNA the protein SMG9 has been proposed to play a role in brain, heart and eye development. STX8 is thought to be involved in vesicle trafficking and POLA1 in the initiation of DNA synthesis. The role of SPAG7 is so far unknown.

**Figure 1:**
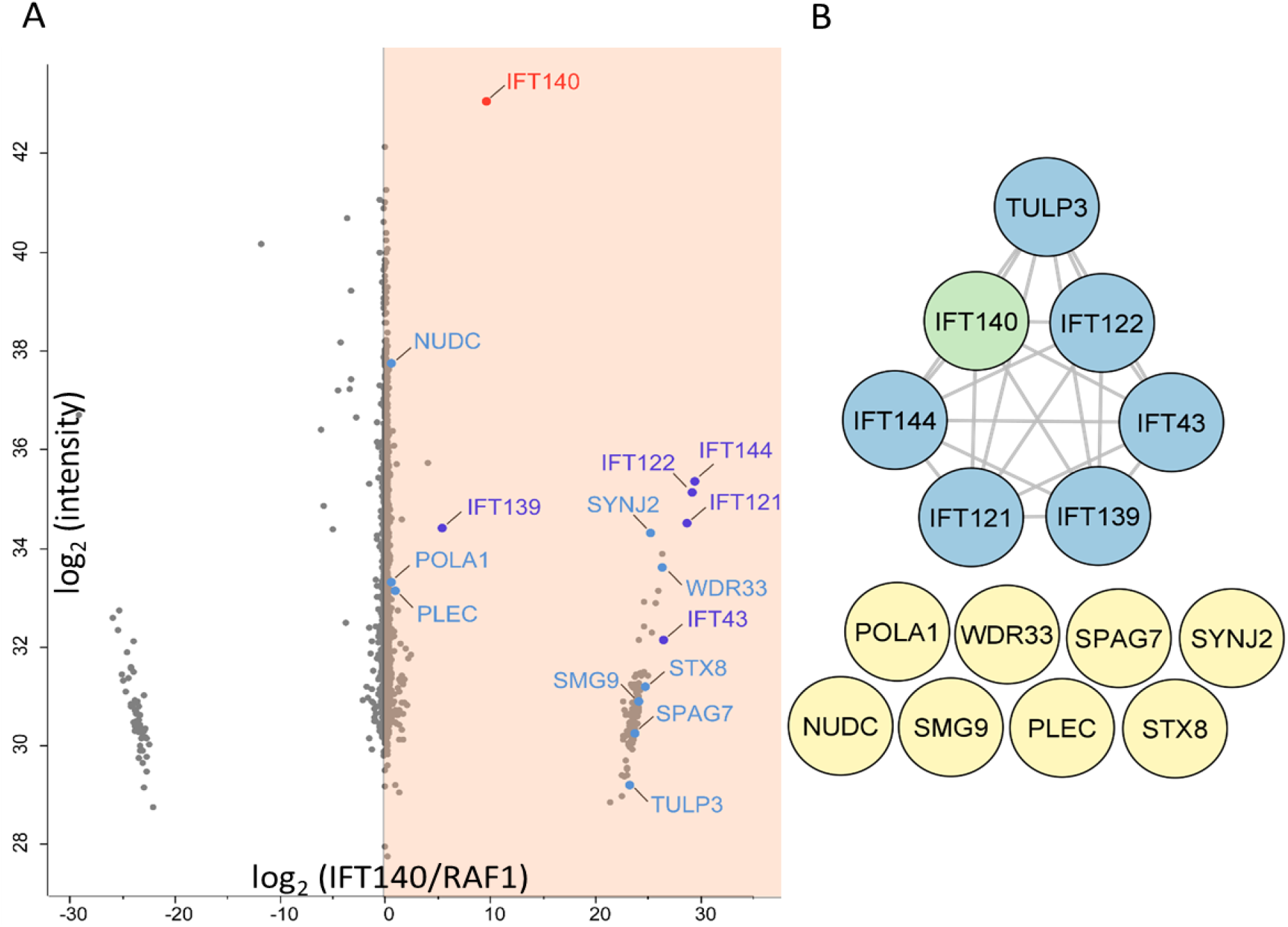
Identification of the IFT140 wildtype interactome. (A) Using Strep immunoprecipitation (IP) with lysates from HEK293 cells stably overexpressing IFT140 WT or RAF1 as negative control fused to an N-terminal Strep/FLAG-tag, we determined the protein interactome of IFT140. Mass spectrometric identification and label free quantification of the detected peptides was performed using MaxQuant. Statistical analysis (n=36) was done with Perseus (t-test: p<0.05; Significance A: p<0.05) and the log_2_ ratios of IFT140 vs. RAF1 calculated and plotted. (B) A protein interaction analysis for the identified interactors was performed using the STRING database. In addition to all components of the IFT-A complex also known interactors like TULP3 were identified, confirming the suitability of our approach to detect interaction partners of IFT140. In total 9 proteins which are not part of the IFT-A complex were identified as potential interactors.

**Table 1:** Enriched proteins in the IFT140 wildtype immunoprecipitation. For the analysis of the IFT140 protein-protein interaction a Strep immunoprecipitation (IP) was performed (n=36). RAF1 was used as a negative control. Mass spectrometric identification and label free quantification of the detected peptides was performed using MaxQuant. Statistical analysis was done using a R script and the log_2_ ratios calculated (IFT140/RAF1). Ratios as well as statistical significance (*=p<0.05; **=p<0.01; ***=p<0.001) from students’s t-test and significance A and known functions of potential interactors are shown.

| Gene name | log<sub>2</sub> ratio (mut/wt) | student's t-test | Significance A | Protein function |
| --- | --- | --- | --- | --- |
| IFT144 | 29.45 | *** | *** | IFT-A core complex (Nakayama and Katoh 2020) |
| IFT121 | 28.73 | *** | *** | IFT-A peripheral complex (Nakayama and Katoh 2020) |
| IFT43 | 26.44 | *** | *** | IFT-A peripheral complex (Nakayama and Katoh 2020) |
| WDR33 | 26.31 | * | *** | mRNA modification |
| SYNJ2 | 25.18 | * | *** | membrane trafficking and signal transduction (Chuang et al. 2012) |
| STX8 | 24.71 | * | *** | vesicle trafficking (Yanguas and Valdivieso 2021) |
| SMG9 | 24.14 | * | *** | Wnt/β-catenin signaling (Jin et al. 2021) |
| IFT122 | 23.71 | *** | *** | IFT-A core complex (Nakayama and Katoh 2020) |
| SPAG7 | 23.71 | ** | *** | unknown |
| TULP3 | 23.18 | *** | *** | negative regulation of Shh-signaling (Hong et al. 2021; Mukhopadhyay et al. 2013) |
| IFT140 (bait) | 9.6 | *** | *** | IFT-A core complex (Nakayama and Katoh 2020) |
| IFT139 | 5.39 | ** | *** | IFT-A peripheral complex (Nakayama and Katoh 2020) |
| PLEC | 0.88 | * | *** | cytoskeleton, interlinks intermediate filaments with microtubules (Kaneyasu et al. 2022) |
| POLA1 | 0.59 | ** | ** | initiation of DNA synthesis (Starokadomskyy et al. 2016) |
| NUDC | 0.53 | *** | ** | Spindle formation and microtubule organization (Biebl et al. 2022; Aumais et al. 2003) |

**Table 2:** Missense mutations in IFT140 and corresponding clinical phenotype. From previous publications we gathered a list of 24 missense mutations reported from patients suffering from ciliopathies and harboring mutations in both alleles of IFT140, mostly compound heterozygotes. For each patient the allelic combination and the reported clinical phenotype is shown. Mutations are sorted according to severity of the clinical phenotype.

| Phenotype | Mutation | Second allele | Publication |
| --- | --- | --- | --- |
| Isolated retinal dystrophy | p. C333Y | p. C333Y | (Hull et al. 2016) |
| Isolated retinal dystrophy | p. A341T | p. Arg475Asnfs*14 | (Hull et al. 2016) |
| Isolated retinal dystrophy | p. T484M | p. T484M | (Hull et al. 2016) |
| Isolated retinal dystrophy | p. S939P | c.2399+1G>T | (Hull et al. 2016) |
| retinitis pigmentosa | p. P71L | p. V217Gfs*2 | (Xu et al. 2015a) |
| retinitis pigmentosa | p. C663W | p. G1276R | (Xu et al. 2015a) |
| retinitis pigmentosa | p. R871C | p. W459* | (Xu et al. 2015a) |
| retinitis pigmentosa | p. A974V | p. A418P | (Xu et al. 2015a) |
| retinitis pigmentosa | p. L1399P | p. N633Sfs*10 | (Xu et al. 2015a) |
| LCA | p. C329R | p. T484M | (Xu et al. 2015a) |
| LCA | p. E790K | p. E522Gfs*6 | (Xu et al. 2015a) |
| Atrioventricular septal defect | p. V108M | p. E1065K | (Priest et al. 2016) |
| LCA, renal failure | p. L440P | unknown | (Hull et al. 2016) |
| Retinal dystrophy with skeletal defects | p. E664K | p. E664K | (Khan, Bolz, and Bergmann 2014b; Miller et al. 2013) |
| JATD | p. G212R | c.2399+1G>T | (Perrault et al. 2012a) |
| JATD | p. V292M | p. G522E | (Schmidts et al. 2013) |
| JATD | p. V292M | p. N460Kfs28 | (Schmidts et al. 2013) |
| Mainzer-Saldino | p. G140R | p. E164* | (Schmidts et al. 2013) |
| Mainzer-Saldino | p. G212R | p. Ala1306Glyfs*56 | (Perrault et al. 2012a) |
| Mainzer-Saldino | p. I233M | c.2399+1G>T | (Perrault et al. 2012a) |
| Mainzer-Saldino | p. I233M | p. I233M | (Perrault et al. 2012a) |
| Mainzer-Saldino | p. Y311C | p. Ile286Lysfs*6 | (Perrault et al. 2012a) |
| Mainzer-Saldino | p. E664K | c.2399+1G>T | (Perrault et al. 2012a) |
| Mainzer-Saldino | p. C1360R | c.2399+1G>T | (Schmidts et al. 2013) |

### Collecting missense mutations in IFT140 from ciliopathy-affected patients

From previous publications we gathered a list of 24 missense mutations in IFT140 found in patients affect by ciliopathies (Table 2). 20 out of 24 of these mutations occur as compound heterozygotes while 4 mutations (I233M, C333Y, T484M and E664K) occur homozygous in patients (Khan, Bolz, and Bergmann 2014a; Xu et al. 2015b; Bifari et al. 2016; Hull et al. 2016; Priest et al. 2016; Peña-Padilla et al. 2017; Schmidts et al. 2013). These 24 mutations are associated with retinal diseases as well as skeletal abnormalities, atrioventricular septal defects and organ deficiencies such as Mainzer-Saldino and Jeune syndrome (see Table 2). In case of the four patients with homozygous mutations in IFT140 the genotype most likely can be more directly correlated to the clinical phenotype observed in the affected patients. For analysis of the PPIs of all 24 mutations we generated HEK293 stable lines expressing IFT140 WT or ciliopathy-related mutant constructs with a N-terminal Strep/FLAG-tag. A structure prediction from alphafold for human IFT140 with the position of each mutation indicated is shown in supplement figure 2.

### Ciliopathy-associated IFT140 missense mutations cause interactome changes

To analyse the IFT140 protein interactome and the changes induced by ciliopathy-associated mutations we performed affinity purification followed by LC-MS/MS with the generated HEK293 stable lines. After cell lysis followed by affinity purification a quantitative LC-MS/MS analysis was performed. Using label free quantification of the potential interaction partners we first filtered for stable interactors of WT IFT140. In a second step the ratio of these interactors was evaluated for quantitative changes in the mutants compared to the WT. Through the statistical analysis of 6 replicates each using stringent statistical filters (p-value <0.05 and Significance A p-value <0.05) we were able to gain robust and reproducible results. As examples, the effects of IFT140 missense mutations on complex stability and TULP3 interaction are shown for the mutations G212R, E267G and C663W in Figure 2. In Figure 3 all results regarding IFT-A and TULP3 interaction are summarized.

**Figure 2:**
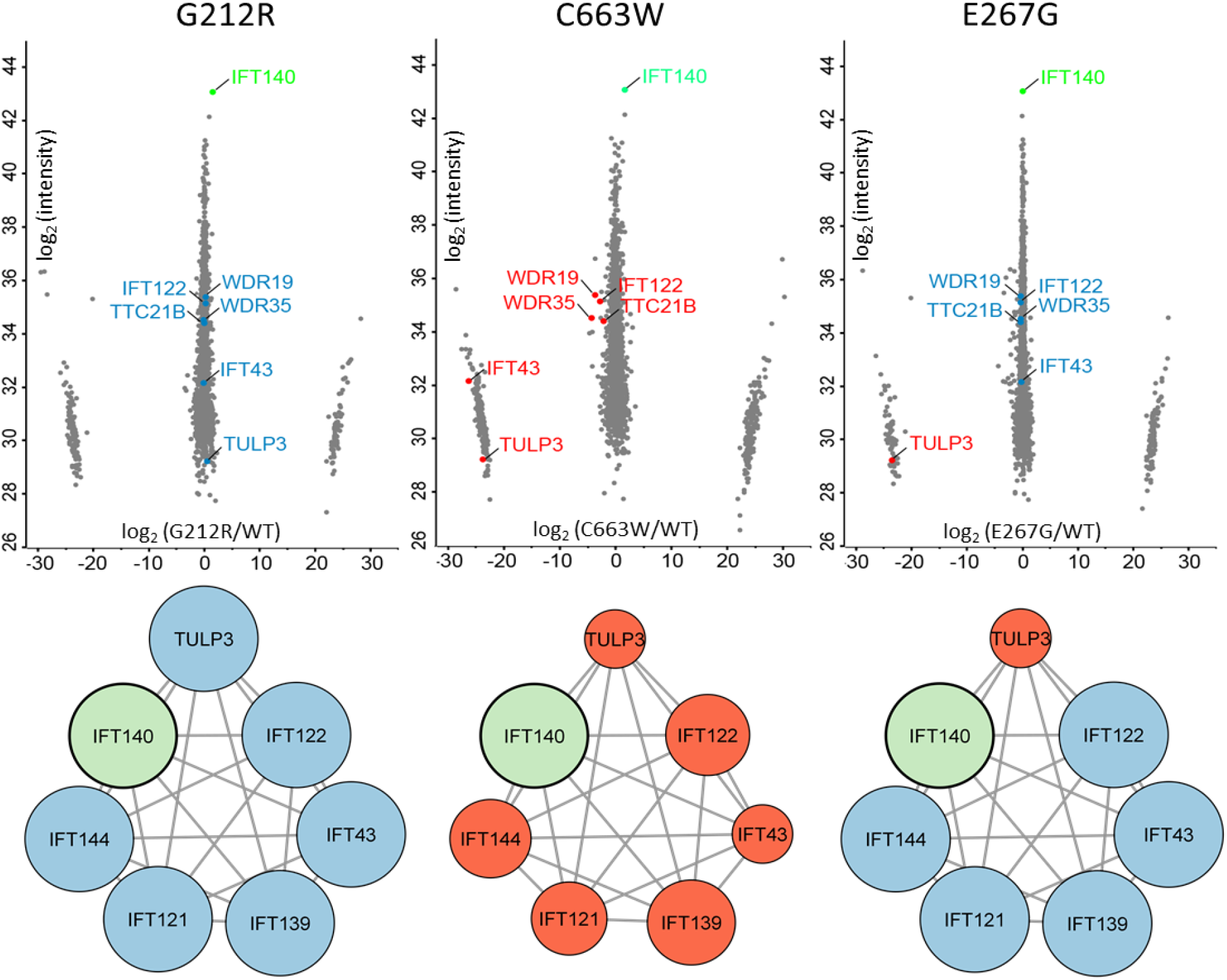
Missense mutations in IFT140 have varying effects on IFT-A complex stability and TULP3 interaction. The effect of missense mutations in IFT140 were assessed via Strep immunoprecipitation (IP) with lysates from HEK293 cells stably overexpressing IFT140 WT or mutant construct fused to an N-terminal Strep/FLAG-tag. Mass spectrometric identification and label free quantification of the detected peptides was performed using MaxQuant. Statistical analysis was performed using a R script and the log_2_ ratios of mutant vs. WT were calculated. Scatter plots are shown for the mutations G212R, E267G and C663W. The effect of each mutation on the interaction of IFT140 with other IFT-A proteins as well as TULP3 is shown below each scatter plot. IFT140 (green) was used as bait. Protein interactions that have not been affected are shown in blue. Disrupted interactions are indicated in red with the size reduction of the corresponding circle reflecting the severity of the abundance reduction in the mutant as compared to IFT140 WT.

**Figure 3:**
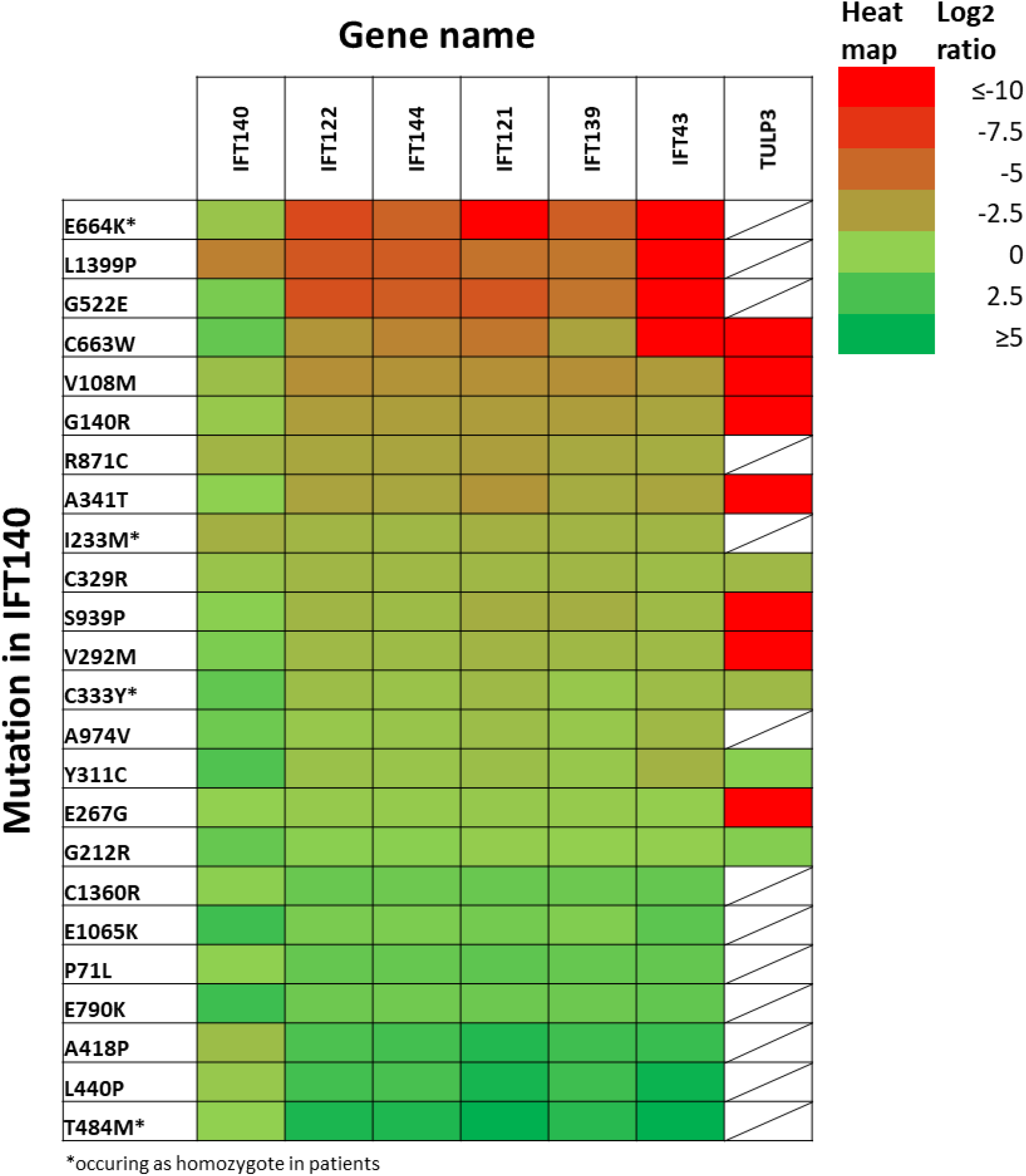
A subset IFT140 missense mutations are hypomorph and have a quantitative effect on IFT-A complex stability. For 24 ciliopathy-associated missense mutations in IFT140 the changes affecting interaction partners of IFT140 WT were assessed by Strep immunoprecipitation. Mass spectrometric identification and label free quantification of the detected peptides was performed using MaxQuant. Statistical analysis was performed using a R script and the log_2_ ratios of mutant vs. WT were calculated and are shown in a heat map for 24 missense mutations. Log_2_ ratios of 0 indicate similar abundance of the protein in WT and mutant conditions, while log_2_ ratios <0 indicate reduced protein levels in mutant samples as compared to WT. White fields are shown for cases in which no data could be obtained. Mutations are sorted from top to bottom according to the severity of the IFT-A complex disruption according to the accumulated negative log_2_ fold change of all IFT-A components.

The IFT-A complex could be reliably detected and quantified in our experiments (Figure 3). For 7 out of 24 mutations (V108M, G140R, A341T, G522E, E664K, C663W, S939P) we could observe an impairment of the IFT-A complex stability (Figure 3, seven mutations on top). This is indicated by the reduced abundance of IFT-A components in these mutants, while the levels of IFT140 are comparable in both conditions. For other mutations the complex composition was not affected (Figure 2 and 3). We therefore assume that at least a subset of missense mutations might disrupt the stability of the IFT-A complex, changing its composition and potentially hampering its molecular function. Several patients harbour two mutations which do not lead to the disruption of the IFT-A complex in our experiments, e.g. A974V/A418P and C329R/T484M. Therefore, disrupted complex stability is unlikely to be the sole cause of disease in these cases. In the mutations A418, L440P and T484M (three out of 24), all components of the IFT-A complex are enriched, even though levels of IFT140 are similar as compared to the WT. In the case of the T484M mutation IFT43 is significantly enriched.

For the mutation L1399P the abundance of the bait protein IFT140 as well as all IFT-A components were significantly reduced in comparison to the WT, indicating problems with expression or protein stability. V108M, G140R, A341T, G522E, E664K, C663W, R871C and L1399P (8 out of 24) led to significantly reduced abundance of IFT-A components. In the case of G522E, E664K, C663W and L1399P (four out of 24) IFT43 could not be detected in the mutant condition.

In the mutant conditions G212R, Y311C, C329R and C333Y (four out of 24), the protein TULP3 was detected at similar levels as in the corresponding WT control. A group of seven out of 24 mutations (V108M, G140R, E267G, V292M, A341T, C663W and S939P) showed a strong reduction of TULP3. We did not identify any cases whereby a higher abundance of TULP3 was observed in the mutant in comparison to the WT condition.

All four mutations (G212R, Y311C, C329R and C333Y) which showed no reduced abundance of TULP3 as compared to IFT140 WT also no strong disruption of IFT-A complex stability was observed (shown for the mutation G212R in Figure 2, Figure 3). In the case of the four mutations (V108M, G140R, A341T and C663W) a severe disruption of IFT-A complex stability and reduced abundance of TULP3 could be detected (shown for the mutation C663W in Figure 2, Figure 3). Three mutations (E267G, V292M and E267G) resulted in reduced TULP3 abundance in the mutant, but had only a weak (V292M and S939P) or no effect at all (E267G) on IFT-A composition (shown for the mutation E267G in Figure 2). Interestingly a group of seven SNPs from IFT140 shows no effect on the integrity of the IFT-A integrity (supplement figure 3)

### TULP3 interaction is impaired to varying degree by IFT140 missense mutations

Our data identified TULP3 as enriched in IFT140 WT, a ciliary protein with a known role in the regulation of hedgehog signalling (Figure 1). In the mutant conditions G212R, Y311C, C329R and C333Y (four out of 24), the protein TULP3 was detected at similar levels as in the corresponding WT control. A group of seven out of 24 mutations (V108M, G140R, E267G, V292M, A341T, C663W and S939P) showed a strong reduction of TULP3 abundance (Figure 2, 3). The observed reduction in TULP3 binding hints towards a potential impairment or even disruptive effect of seven out of 24 of the analysed missense mutations on the interaction between IFT140 and TULP3. For the other 13 out of 24 mutations the data did not allow reliable quantification of TULP3. This might be due to technical limitations of the applied shotgun proteomics approach regarding the detection of lowly abundant proteins in complex samples, which may be the case with TULP3. Even though not all mutants were affected, a misregulation of IFT140-TULP3 interaction to varying degree might be a cause of clinical defects observed in patients. This also holds true when considering the compound heterozygosity of most patients, where the combination of different alleles might cause varying degrees of IFT140-TULP3 disruption. Notably for three of the four mutations (Y311C, C333Y, T484M, E664K) which occur as homozygotes, TULP3 was not stably detected, which led us to targeted investigation of TULP3 interaction using co-immunoprecipitation.

We further examined the interaction of IFT140 and TULP3 using co-immunoprecipitation followed by SDS-PAGE and western blotting using HEK293 cells stably expressing N-terminally SF-tagged IFT140 constructs. We co-transfected these cells with a vector expressing TULP3 fused to a N-terminal HA-tag followed by FLAG affinity purification of IFT140 and detection of IFT140 and TULP3 in eluates by using specific antibodies (Figure 4 and 5).

**Figure 4:**
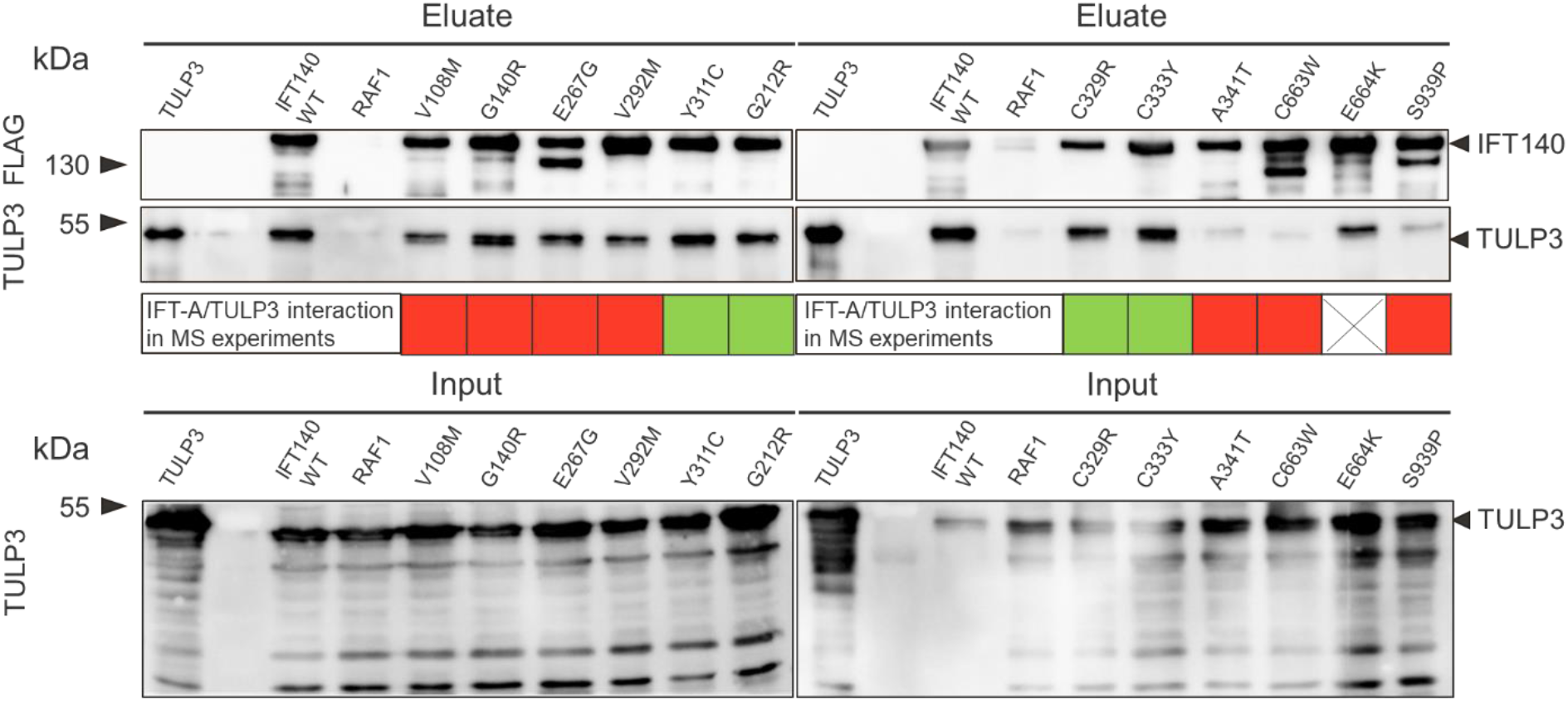
The interaction of IFT140 and TULP3 is disturbed by a subset of missense mutation. For 12 out of 24 mutations (including all for which a mutant/WT ratio for TULP3 could be determined) a co-immunoprecipitation was performed. Following a FLAG immunoprecipitation of (N)SF-tagged IFT140 WT and mutant expressing HEK293 cells transiently co-transfected with (N)HA-TULP3. A SDS-PAGE was performed followed by western blotting with FLAG-specific (for IFT140) and-TULP3-specific antibody. The levels of TULP3 in equal protein amounts of lysate (50ug) are shown as input and indicate comparable levels of TULP3 in lysates. For each analyzed mutant the result for the IFT-A/TULP3 interaction from the MS-experiments is shown at the bottom (red box= disruption of interaction; green box= no disruption of interaction).

**Figure 5:**
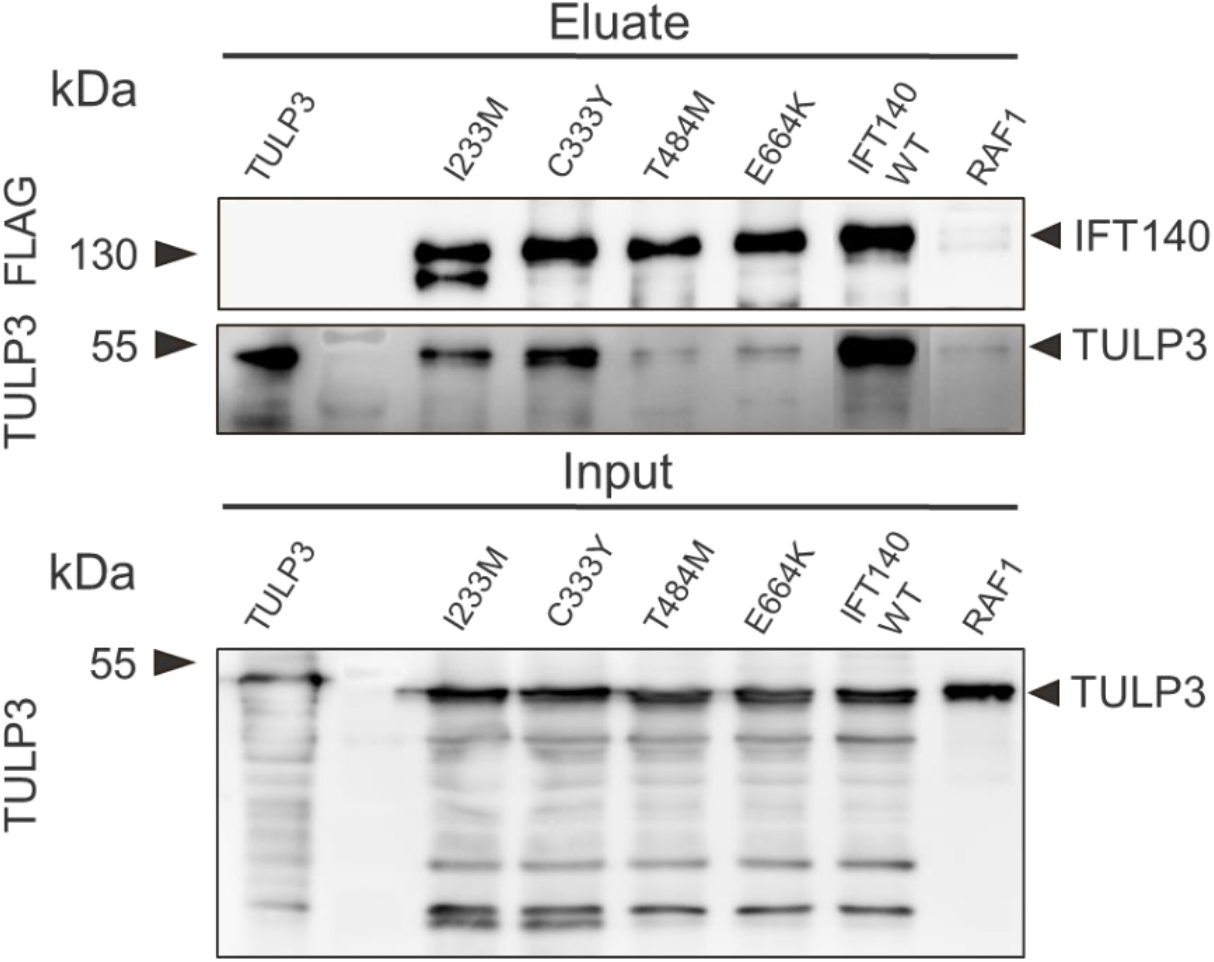
The interaction of IFT140 and TULP3 differs in homozygous missense mutations. For all four mutations (I233M, C333Y, T484M, E664K) which occur as homozygotes a co-immunoprecipitation was performed. Following a FLAG immunoprecipitation of (N)SF-tagged IFT140 Wildtype and mutant expressing HEK293 cells transiently co-transfected with (N)HA-TULP3. A SDS-PAGE was performed followed by western blotting with FLAG-specific (for IFT140) and-TULP3-specific antibody. The levels of TULP3 in equal protein amounts of lysate (50ug) are shown as input and indicate comparable levels of TULP3 in lysates.

The results of the co-immunoprecipitation further confirmed our findings from the interactome studies. The six out of seven mutations (V108M, E267G, V292M, A341T, C663W, S939P) which showed a loss of TULP3 in the previous affinity purification experiments also exhibit reduced binding of TULP3 to IFT140 in co-immunoprecipitation (Figure 4). This effect is not due to different amounts of TULP3, as shown by similar amounts of the protein lysate input. The only exception is the G140R mutation, where the reduced abundance of TULP3 in the proteomics experiments is not reflected by the co-immunoprecipitation. While the amounts of IFT140 in eluates seemed to vary slightly and degradation products were seen in some conditions, these do not contradict our statement. The observed alterations in levels of TULP3 are much stronger than the differences in the abundance of bait protein in eluates. For the E664K mutation no mass spectrometric data was available.

To better understand genotype-phenotype correlations regarding TULP3 a focused analysis of the four homozygous mutations I233M, C333Y, T484M, and E664K was performed (Figure 5). In our proteomics experiment we could only obtain data for C333Y, where the interaction of IFT140 and TULP3 was not changed significantly in comparison to IFT140 WT. In the co-immunoprecipitation a strong signal for TULP3 is visible in case of the C333Y mutation. However, the signal appears weaker in comparison to the WT. In the case of the I233M mutation a degradation product of IFT140 as well as reduced levels of TULP3 can be seen. For T484M and E664K only a weak signal for TULP3 is visible, indicating a strong disruption. Taken together our data strongly suggests that the interaction between IFT140 and TULP3 is disrupted to varying degree across a subset of missense mutations.

## Discussion

In this study we examined the changes in protein interaction networks induced by missense mutations using a proteomics approach. This approach created valuable new insights into the interaction network of IFT140 and the effect of disease-causing mutations on protein-protein interactions (PPIs). By gaining insights into the effects of ciliopathy-associated missense mutations in IFT140 we aimed at better understanding the general mechanisms of genetic disorders on the molecular level leading to a varying degree of disease severity. Understanding the relationship between genotype and phenotype and disease mechanisms causing disease is a major challenge in the understanding of genetic disease and not limited to ciliopathies. Potential mechanisms include e.g. reduced abundance of functional protein due to protein stability, disruption of protein complexes as well as the alteration of specific PPIs. Due to the necessity of functional IFT140 during development it seems unlikely that ciliopathy-associated missense mutations on both alleles of a patients lead to a full loss of protein or function but rather to a limited extent (hypomorph).

### Analysis of IFT140 WT reveals potential new interactors

By analysing a large number of biological replicates (n=36) we were able to obtain crucial and convincing insights in the interaction network of IFT140. The statistical analysis of the data from the label free quantification identified all components of the IFT-A complex as significantly enriched. In addition, known interactors of IFT-A like the ciliary proteins TULP3 and NUDC were enriched in IFT140 IPs. Taken together this data supports the robustness of our approach to reliably identify the interactome of IFT140. An analysis of the interaction landscape using the STRING database also revealed that our dataset contains seven so far unknown potential interactors of IFT140. Among these is the SYNJ2 which was proposed to play a role in membrane trafficking and signal transduction (Chuang et al. 2012). The association of SYNJ2 with the IFT-A complex might hint towards a role of SYNY2 in the trafficking of membrane bound cargo in a ciliary context. This might explain the previously proposed role of SYNJ2 in signal transduction. Another interesting new interactor of IFT140 is SMG9. It has been implicated in brain, heart and eye development (Jin et al. 2021). This is in line with the defects observed in patients suffering from ciliopathies associated to mutations in IFT140 (table 2). The exact role of the interaction between IFT140 and SMG9 and its potential role in ciliary function needs to be further evaluated in the future. NUDC was found as slightly enriched in IFT140 WT and the interaction remains unaltered by missense mutations (supplement figure 1). NUDC is important for mitotic spindle formation and chromosome separation during mitosis (Chen et al. 2015). Members of the nud gene family are essential for brain cortex development and is known to serve as a co-chaperone in protein folding (Zheng et al. 2012; Biebl et al. 2022). NUDC might be important for correct folding of IFT140, especially the WD40 domains, to which it is known to bind (Chen et al. 2015). SPAG7 was also identified as potential interactor of IFT140 and reduced in the case of E790K only. The function of SPAG7 is so far unknown. POLA1 is involved in the initiation of DNA synthesis and was unaffected in all mutant conditions (supplement figure 1). It importance for ciliary function needs to be examined further. PLEC was detected in all conditions and enriched in the case of P71L, L440P, T484M and C1360R (four out of 24) while being mildly reduced in the case of G522E (one out of 24). It is of known importance for interlinking intermediate filaments with microtubules (Wiche 1998). It might be important for formation of ciliary structures and their stability and require IFT140 for correct function and localization which could be investigated in the future.

According to the IntAct database (https://www.ebi.ac.uk/intact) 26 interactors of IFT140 have been reported so far. 14 of these were not found as enriched in our dataset. One reasons for this may be the statistical analysis used in our approach, which leads to a stringent filtering of potential interactors. While excluding other potentially interesting candidates, we generated a list of bona fide interactors (table 1).

### Missense mutations in IFT140 have a quantitative effect on IFT-A complex stability and show a genotype-phenotype correlation

Our mass spectrometric based analysis revealed that ciliopathy-associated missense mutations in IFT140 have a quantitative effect on IFT-A complex stability. Rather than causing a complete disruption of the protein complex a hypomorph effect was observed resulting in reduced native IFT140 based IFT-A complex binding. Seven out of 24 of the analysed mutations (V108M, G140R, A341T, G522E, E664K, C663W, S939P) impair the stability of the IFT-A complex, leading to a partial disruption. Due to the importance of functional IFT-A for retrograde trafficking of cargo towards the basal body (Pazour and Rosenbaum 2002; Hirano et al. 2017; Pazour, Wilkerson, and Witman 1998; Nakayama and Katoh 2020) the disruption of the complex might explain some of the defects observed in patients harbouring these mutations.

Interestingly we can observe a correlation between genotype and phenotype in the eight patients carrying missense mutations on both alleles of IFT140. Patients who harbour two mildly or undisruptive mutations on both alleles show the mildest clinical phenotypes (see table 2, supplement table 1). The disease severity is increased when one allele harbours a missense mutation, which shows strong disruption of IFT-A complex stability in our experiments. Patients with IFT-A disruptive missense mutations on both alleles show the strongest phenotype. This observation is evident of a clear correlation between the severity of PPI disruption observed in our experiments and the disease severity observed in affected patients. This correlation is further strengthened by the fact that six examined single nucleotide polymorphisms (SNPs) in IFT140 do not show any impact on the IFT-A complex stability (supplement Figure 3).

It also indicates that the combination of missense mutations affects the phenotype, as seen in the case of the T484M mutation. While a patient carrying a non-disruptive biallelic T484M mutation shows isolated retinal dystrophy, a patient harbouring the same T484M mutation in combination with the disruptive C329R mutation was diagnosed with LCA. This more severe phenotype, which is characterized by early-onset blindness, may be due to the more severe allelic combination, indicating genetic threshold effects that define disease severity.

### A subset of IFT140 missense mutations shows edgetic effects on TULP3

The potential effect of amino acid changes on protein-protein interactions is known and has been observed for a broad variety of proteins (Boldt et al. 2016; Del-Toro et al. 2019). One model to explain these alterations is the edgetic concept, in which mutations lead to the loss of a specific PPI (edge loss) rather than to a total loss of function of the affected protein (node loss) (Zhong et al. 2009; Wang, Sahni, and Vidal 2015). Our data shows that mutations in IFT140 can impair specific interactions within an interaction network, for example TULP3. This is consistent with the edgetic hypothesis and further strengthens the idea of edge removal in protein interaction networks induced by missense mutations.

Interestingly, we also observed a disruption of the interaction between IFT-A and TULP3 caused by seven out of 24 mutations (V108M, G140R, E267G, V292M, A341T, C663W, S939P) in IFT140 in MS/MS-experiments. For four out of 24 mutations (G212R, Y311C, C329R, C333Y) the interaction with TULP3 was slightly reduced or remained unchanged in comparison to the WT. For the other 13 out of 24 mutations no ratio for TULP3 could be obtained. We further validated the interaction of IFT140 and TULP3 via co-immunoprecipitation for 15 out of 25 mutations, including all homozygotes and all mutations for which a TULP3 ratio could be obtained in the proteomics experiment. The degree of the loss of TULP3 varied between mutations in co-immunoprecipitation experiments (Figure 4 and 5). This indicates that the loss is not complete, but rather that mutations in IFT140 reduced the interaction with TULP3 to varying degrees. For the homozygous mutations a weak disruption of the interaction with TULP3 could be observed for C333Y and to a stronger extent for I233M. For the homozygous mutations T484M and E664K a stronger disruption was observed in the interaction between IFT140 and TULP3. Keeping in mind the importance of the IFT140/TULP3 interaction during development, the question remains open which level of TULP3 interaction results in reduction or even loss of signalling, or if other IFT-A complex proteins might compensate some function. Interestingly T484M, resulting in a mild phenotype, shows normal IFT-A binding, whereas E664K showed loss of IFT-A and TULP3, resulted in additional skeletal defects. However, due to the low number of patients, it should be validated, to what extend TULP3 dependent signalling is affected by specific mutations. In addition, it remains unknown if this effect is also true *in vivo*, which should be examined in future experiments. Knockout of TULP3 in mice leads to disturbed eye development, neural tube defects and other developmental defects (Norman et al. 2009; Ikeda et al. 2001), which shows an clinical overlap with some of the observed defects in patients suffering from IFT140-associated ciliopathies. In hTERT-RPE1 cells knockout of TULP3 leads to defective cilia formation (Hong et al. 2021; Han et al. 2019) and it could be shown that TULP3 bridges the IFT-A complex to the ciliary membrane, which promotes trafficking of components important for hedgehog signaling, such as Gpr161, a negative regulator of hedgehog signalling pathway (Mukhopadhyay et al. 2010). In cells lacking TULP3 in their primary cilia due to knockout of IFT-A components also Gpr161 was absent from the cilium (Hirano et al. 2017). Primary cilia are essential for functional hedgehog signalling. Defects in hedgehog signaling have been implicated in several ciliopathies, including retinal dystrophies as well as skeletal abnormalities and organ damage (Anvarian et al. 2019; Waters and Beales 2011; Lai et al. 2011). The ciliary dislocation of these proteins caused by the disrupted IFT140/TULP3 interaction might provide a possible explanation how IFT140 defects lead to skeletal abnormalities and organ defects in patients.

## Conclusion

Taken together our data indicates that disease-associated missense mutations in IFT140 are hypomorph and have edgetic effects. The genotype severity of the IFT-A complex disruption observed in our experiment also allows us to establish a correlation between genotype and clinical phenotype. The disruption of the stability of the IFT-A complex seen for hypomorph mutations in IFT140 occurs in a quantitative manner. This might hamper the complex and limit its ability to function properly, potentially leading to transport defects resulting in mislocalization of important cargo for proper ciliary function. The disruption of specific PPIs, as shown here e.g. for TULP3, further strengthens the idea of edgetic perturbations caused by disease-associated mutations. It also opens the path to link mutations to specific disease mechanisms like hedgehog signaling. The disruption of different PPIs to a varying degree as well as the allelic combination also might explain the clinical variation observed in patients regarding type, severity and tissue specificity of the observed ciliopathies. Since our experiments were conducted in HEK293 cells these finding need to be further validated in the future in a more suitable model system, e.g. patient-derived cells or *in-vivo* mouse models.

Our findings do not allow to propose a disease mechanism for each of the observed mutations or patients. This further highlights the existing of additional effects, which have not been identified so far. One possibility might be threshold effects caused by altered levels of functional protein, which could include folding defects caused by mutations, leading to the degradation of large amounts of IFT140 in the cell or the accumulation of misfolded protein with limited functionality. In addition, it seems likely that, at least for some of the existing disease mechanisms, compensatory effects or modifiers are relevant, e.g. chaperones. Additional research is necessary to further illuminate these interplays.

## Acknowledgements

This work was funded by Wellcome Trust (210585/A/18/Z) and in part the Kerstan Foundation to Prof. Marius Ueffing.

Thanks to the groups of Johannes Gloeckner and Ronald Roepman for kindly supplying plasmids.

Thanks to Felix Hoffmann and Shibu Antony for scientific input, Mohamed Ali Jarboui for helping with the data analysis and Christine Henes for helping with experiments.

## Experimental Procedures

### Site-directed mutagenesis

A wildtype IFT140 entry-construct for gateway cloning was generated from cDNA as template for PCR. Mutant constructs were generated using primers carrying the desired mismatch to obtain the specific mutation. Following PCR a Dpn1 digest was performed for 1h to digest methylated template DNA. The constructs were transformed into DHC5α cells (Invitrogen, USA) and bacteria grown in kanamycin selection conditions. DNA was isolated using a PureYield Plasmid Midi Preparation Kit (Promega, USA) and clones verified via sequencing. A gateway LR reaction with a destination vector harbouring a N-terminal Strep/FLAG-tag was performed, constructs transformed into DHC5α cells (Invitrogen, USA) and bacteria grown in ampicillin selection conditions prior to DNA isolation as described above. All expression constructs were verified via sequencing

**Table 1:**
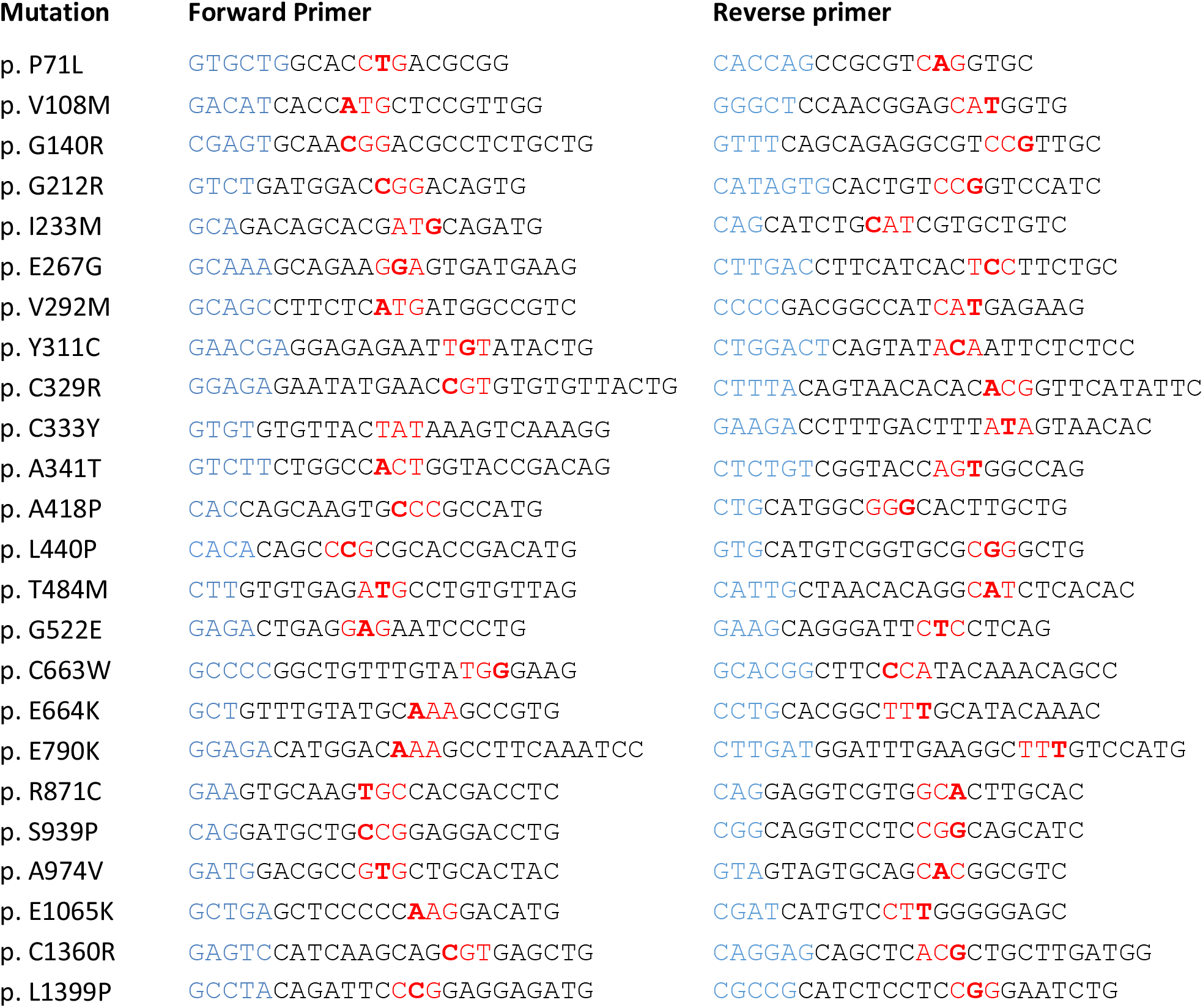
DNA oligomers used for generation of missense mutation constructs. Site-directed mutagenesis primers for each missense mutation. Overhangs are highlighted in blue, base pairs coding for the targeted amino acid change in red with the exact position of the missense mutation indicated in bold letters.

### Plasmids

For Gateway cloning a pDONR201 vector with a Kanamycin resistance (Invitrogen, USA) and pDEST vector with a pcDNA 3.0 backbone and Ampicillin resistance (Invitrogen, USA; modified with N-terminal Strep/FLAG (Gloeckner, Boldt, and Ueffing 2009) or HA tag) were used.

### Generation of stable cell lines

HEK293 (CRL-1573-ATCC) were grown in DMEM (Sigma-Aldrich, USA) supplemented with 10 % FBS (Sigma-Aldrich, USA) and 0.5 % Penicillin/Streptomycin at 37°C and 5 % CO_2_. Cells were transfected using homemade polyethylenimine (PEI) solution and selected for 3 weeks in 500µM Neomycin (Carl Roth, Germany) condition for cells that stably integrated the expression construct. Cells were regularly checked for mycoplasma by PCR.

### Protein complex analysis by Streptavidin affinity purification

For interactome identification of IFT140 WT and mutants, HEK293 control and N-terminal Strep/Flag-tagged IFT140 construct expressing stable HEK293 cells were used with Strep/Flag tagged RAF1 serving as negative control. Cells were cultured as described above and grown in 14 cm plates to 90 %-100 % confluency. Cells were lysed with 1 ml lysis buffer (0.5 % Nonident-P40, 2 % protease inhibitor cocktail (Roche, Switzerland) and 1 % phosphatase inhibitor cocktails I and II (Sigma-Aldrich, USA) in TBS (30 mM Tris-HCl, pH 7.4 and 150 mM NaCl) by rotating for 20 min at 35 rpm at 4°C. After centrifugation for 10 min at 10,000 g at 4°C the protein concentration was measured using Bradford assay and an identical protein amount (5 mg) for each sample was incubated with Strep-Tactin Superflow beads (IBA, Germany) for 1h at 4°C while rotating at 35 rpm. The beads were washed three time with washing buffer (0.1% Nonident-P40 and 1% phosphatase inhibitor cocktails I and II (Sigma-Aldrich, USA) in TBS) and bound proteins eluted with Strep elution buffer (IBA, Germany). An acetone-based protein precipitation was performed followed by tryptic digest and desalting of digested proteins via stop-and-go extraction tips (Thermo Fisher Scientific, USA) as described (Beyer et al. 2018) and prepared for mass spectrometry (MS) analysis.

### Co-immunoprecipitation by FLAG affinity purification

For validation of specific PPIs HEK293 cells stably expressing Strep/Flag-tagged IFT140 or RAF1 constructs were co-transfected with N-terminal HA-tagged TULP3 using homemade PEI solution. After lysis as described above, the protein concentration was measured using Bradford assay (Bio-Rad, USA) and an identical protein amount (2 mg) for each sample was incubated with anti-FLAG M2 agarose beads (IBA, Germany) for 1 hour followed by three washing steps ((0.1 % Nonident-P40 and 1% phosphatase inhibitor cocktails I and II (Sigma-Aldrich, USA) in TBS) and elution of bound protein with FLAG peptide in TBS.

### SDS-PAGE and Western Blot

For investigation of IFT proteins and TULP3, SDS-PAGE using a 10 % Tris-Glycine based separation gel and running buffer was performed with generated eluates from co-immunoprecipitation. Full wet tank blotting (Bio-Rad, USA) was followed by 5 % milk block as well as primary antibodies FLAG-M2-Peroxidase (1:2500, Sigma-Aldrich, USA), TULP3 (1:100; Proteintech, Germany) overnight and secondary goat α rabbit antibody (1:10,000; Jackson ImmunoResearch, USA) for 1h at RT. Membranes were treated with ECL (Thermo Fisher Scientific, USA) and images were taken using the Fusion FX7 imaging system (Vilber, France).

### Experimental Design and Statistical Rationale

For LC-MS/MS analysis an Ultimate3000 nano-RSLC (Thermo Scientific) was coupled to a Orbitrap Fusion Tribrid (Thermo Scientific) by a nanospray ion source. Prepared peptide mixtures were loaded automatically onto a nano trap column (µ-Precolumn 300µm i.d. x 5 mm, packed with Acclaim PepMap100 C18, 5 µm, 100 Å; Dionex, USA). Injection was conducted with a flow rate of 30 µL/min in 98 % of buffer C (0,1 % TFA in HPLC-grade water) and 2 % of buffer B (80 % ACN, 0.08 % formic acid in HPLC-grade water). After 3 min peptides were eluted and separated on an analytical column (75 µm × 25 cm, packed with Acclaim PepMap RSLC, 2 µm, 100 Å; Dionex, USA) at a flow rate of 300 nL/min with a linear gradient from 2 % up to 30 % of buffer B in buffer A (2 % ACN, 0.1 % formic acid) for 82 min after an initial step of 3 min at 2 % buffer B. Remaining peptides were eluted with a steep gradient (30 % to 95 % in 5 min) followed by 5 min at constant 95 % of buffer B before the gradient was decreased rapidly in 5 min to 2 % of solvent B for the final 20 minutes. In the data-dependent analysis full scan MS spectra were measured on the Fusion in a mass-charge range from m/z 335-1,500 with a resolution of 70,000. The ten most abundant precursor ions were selected with a quadrupole mass filter, if they exceeded an intensity threshold of 5.0 x10^4^ and were at least doubly charged, for further fragmentation using higher energy collisional dissociation (HCD) set at a value of 30 followed by mass analysis of the fragments in the iontrap at a resolution of 120,000. Lock mass option was activated and the background signal with a mass of 445.12003 was used as mass lock. The selected ions were excluded for further fragmentation the following 20 s.

MaxQuant software 1.6.1.0 (Cox and Mann 2008) was used for label free quantification. Trypsin/P was used as a digestion enzyme with a maximum of 2 missed cleavages. Cysteine carbamidomethylation was set as fixed modification and methionine oxidations as well as N-terminal acetylation as variable modifications. Minimum ratio count was set at 2 and the first search peptide tolerance at 20. Main search peptide tolerance was set at 4.5 ppm and the re-quantify option was selected. Current SwissProt database (release 2019_11) was used and contaminants detected by the MaxQuant contaminant search. A minimum of two peptides with at least one unique and one razor peptide was required with a minimum length of 7 amino acids for quantification. Match between runs was performed with a match time window of 0.7 min and an alignment window of 20 min.

Statistical analysis was performed using a custom-made R-script. A minimum of 6 biological replicates grown on individual seeded 14 cm dishes were used for statistics, including IFT140 wildtype and RAF1 control during each individual experiment. Data was filtered for potential contaminants and peptides only identified by site and reverse database. For each gene comparison between mutants/RAF1 versus wildtype, at least half of the samples within one group needed to have valid values i.e. detected by the mass spectrometer. The means were compared using a Student’s t-test and p-values were corrected using the Benjamini-Hochberg method (q-values). The log_2_ ratios of each comparison were used to carry out the outlier significance A test. Proteins with a q <= 0.05 or a significance A value <= 0.05 were defined as significantly enriched/depleted.

