## Supplement for "Ciliopathy-associated missense mutations in IFT140 are hypomorphic and have edgetic effects on protein interaction networks"

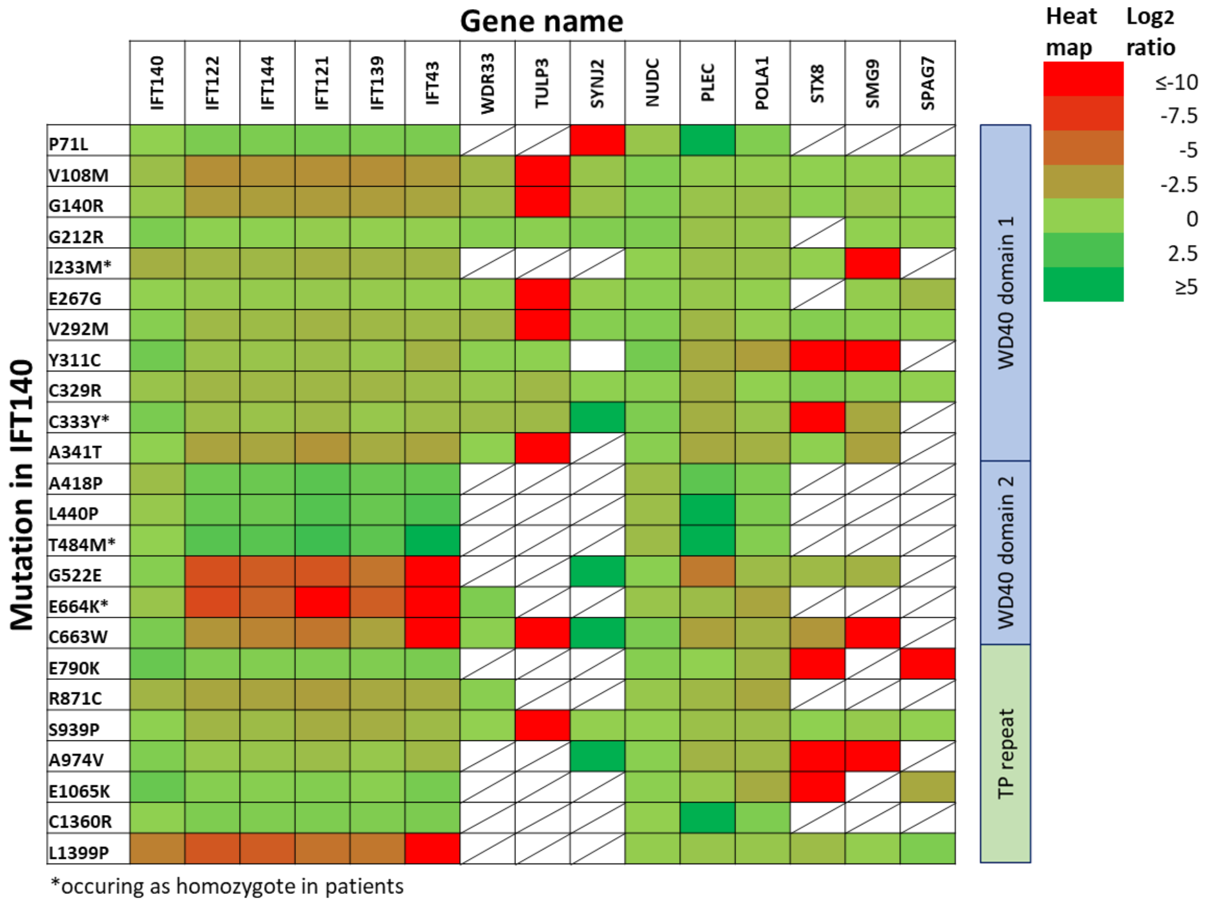


***Supplement figure 1: IFT140 missense mutations have edgetic effects on protein interaction networks***. *For 24 ciliopathy-associated missense mutations in IFT140 the changes affecting interaction partners of wildtype IFT140 were assessed. We applied Step immunoprecipitation (IP) with lysates from HEK293 cells stably overexpressing IFT140 wildtype or mutant construct fused to an N-terminal Strep/FLAG-tag. Mass spectrometric identification and label free quantification of the detected peptides was performed using MaxQuant. Statistical analysis was performed using a R script and the log2 ratios of mutant vs. wildtype calculated and are shown in a heat map for 24 missense mutations. Mutations which occur as homozygotes in patients are indicated in green. Log2 ratios of 0 indicate similar abundance of the protein in wildtype and mutant conditions, while log2 ratios <0 indicate reduced protein levels in mutant samples as compared to wildtype. Crossed fields are shown for cases in which no data could be obtained. Mutations are sorted from top to bottom according to their position in the protein from N- to C-terminal direction*

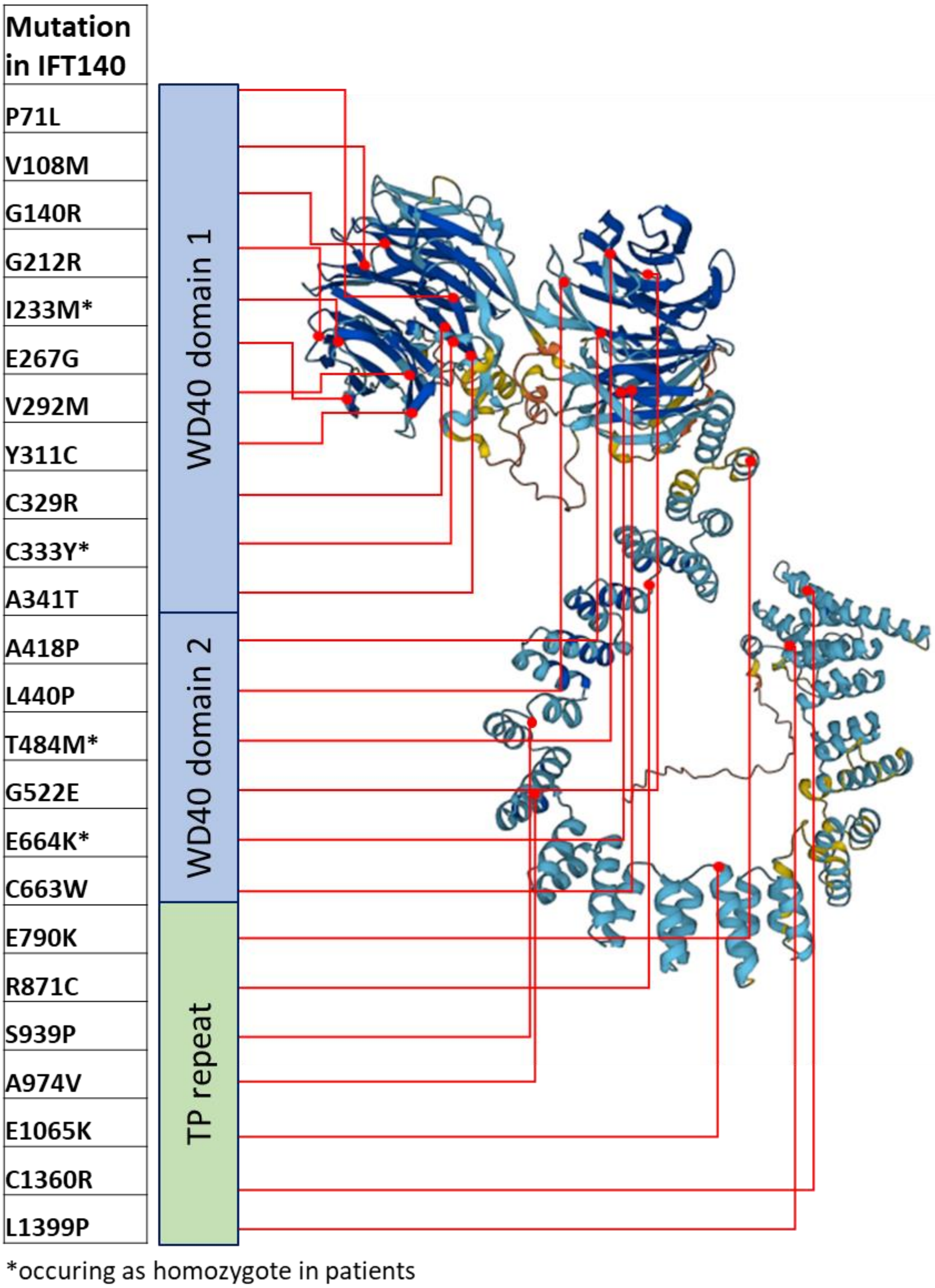


***Supplement figure 2: Position of IFT140 missense mutations in the structure of human IFT140**. Using alphafold a structure prediction for human IFT140 was obtained. The position of each of the 24 missense mutations used in this study is indicated.*

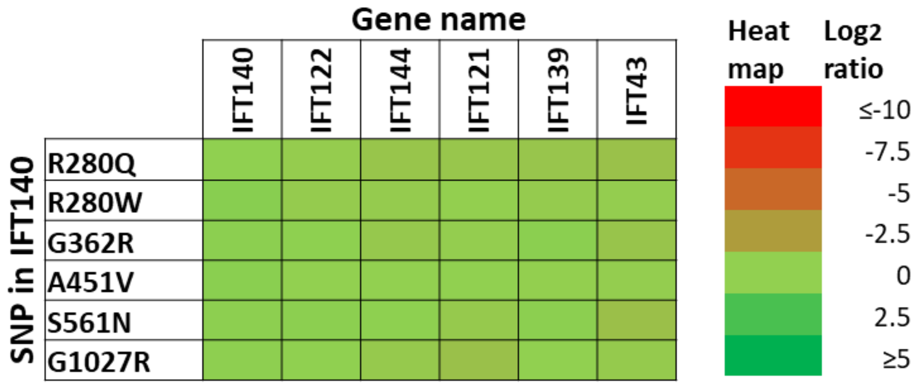


***Supplement figure 3: IFT140 SNPs have no effect on IFT-A complex stability.*** *For six single nucleotide polymorphisms (SNPs) that lead to the exchange of a single amino acid in IFT140 the effect on IFT-A complex stability was assessed. Applying Strep immunoprecipitation followed by mass spectrometric identification and label free quantification of the detected peptides using MaxQuant IFT140 WT and SNPs were compared. Statistical analysis was performed using a R script and the $log_2$ ratios of mutant vs. WT were calculated and are shown in a heat map for six SNPs. $Log_2$ ratios of 0 indicate similar abundance of the protein in WT and mutant conditions, while $log_2$ ratios <0 indicate reduced protein levels in mutant samples as compared to WT.*

***Supplement table 1: Allelic combination in patients reveals correlation between genotype severity and clinical phenotype.*** *For 8 patients who harbor missense mutations on both alleles we assessed the severity of the disruption of IFT-A complex stability (IFT-A integrity) for each allele. The IFT-A integrity is measured by the mean of the $log_2$-change of all IFT-A components in the MS/MS-experiments. A strong disruption is indicated in red, while weak or no disruption is indicated by a green field. Patients are sorted from top to bottom with increasing severity of the clinical phenotype. While patients with two undisruptive or mild disruptive alleles show milder clinical phenotypes, disease severity increases in cases where one strongly disruptive allele is included. The strongest clinical phenotypes are observed in patients who harbor disruptive missense mutations in IFT140 on both alleles, showing a clear correlation between genotype and phenotype.*

| **Patient** | **First allele** | **IFT-A integrity** | **Second allele** | **IFT-A integrity** | **Phenotype** |
|---|---|---|---|---|---|
| 1 | C333Y | -0.38 | C333Y | -0.38 | Isolated retinal dystrophy |
| 2 | T484M | -0.12 | T484M | -0.12 | Isolated retinal dystrophy |
| 3 | A418P | 0.10 | A974V | 0.30 | retinitis pigmentosa |
| 4 | C329R | -1.05 | T484M | -0.12 | LCA |
| 5 | V108M | -2.59 | E1065K | -0.28 | Atrioventricular septal defect |
| 6 | E664K | -12.34 | E664K | -12.34 | Retinal dystrophy with skeletal defects |
| 7 | V292M | -0.80 | G522E | -7.95 | JATD |
| 8 | I233M | -1.35 | I233M | -1.35 | Mainzer-Saldino |

**Supplement table 2: IFT-A complex integrity in IFT-A missense mutations and SNPs.** For 24 disease-associated missense mutations and 8 SNPs in IFT140 we assessed the IFT-A integrity. The IFT-A integrity is measured by the mean of the $log_2$-change of all IFT-A components in the MS/MS-experiments for the corresponding mutation

| **Mutation** | **IFT-A integrity** |
|---|---|
| E664K* | -12.34 |
| L1399P | -8.41 |
| G522E | -7.95 |
| C663W | -6.26 |
| V108M | -2.59 |
| G140R | -2.03 |
| R871C | -1.89 |
| A341T | -1.76 |
| I233M* | -1.35 |
| C329R | -1.05 |
| S939P | -1.04 |
| V292M | -0.80 |
| C333Y* | -0.38 |
| P71L | -0.36 |
| *R280W* | -0.35 |
| E1065K | -0.28 |
| *R280Q* | -0.22 |
| *G1027R* | -0.12 |
| E790K | 0.00 |
| L440P | 0.04 |
| E267G | 0.08 |
| A418P | 0.10 |
| A974V | 0.30 |
| C1360R | 1.19 |
| *G362R* | 1.33 |
| *S561N* | 1.36 |
| G212R | 1.59 |
| *A451V* | 2.35 |
| Y311C | 2.79 |
| T484M* | 7.23 |

***Table 3: Missense mutations in IFT140 and corresponding clinical phenotype***. *From previous publications we gathered a list of 24 missense mutations reported from patients suffering from ciliopathies and harboring mutations in both alleles of IFT140, mostly compound heterozygotes. For each patient the allelic combination, the IFT-A integrity from the MS/MS-experiments and the reported clinical phenotype is shown. Mutations are sorted according to severity of the clinical phenotype.*

| **Phenotype** | **Mutation** | **IFT-A integrity** | **Second allele** | **Publication** |
|---|---|---|---|---|
| Isolated retinal dystrophy | p. C333Y | -0.38 | p. C333Y | (Hull et al. 2016) |
| Isolated retinal dystrophy | p. A341T | -1.76 | p. Arg475Asnfs*14 | (Hull et al. 2016) |
| Isolated retinal dystrophy | p. T484M | 7.23 | p. T484M | (Hull et al. 2016) |
| Isolated retinal dystrophy | p. S939P | -1.04 | c.2399+1G>T | (Hull et al. 2016) |
| retinitis pigmentosa | p. P71L | -0.36 | p. V217Gfs*2 | (Xu et al. 2015) |
| retinitis pigmentosa | p. C663W | -6.26 | p. G1276R | (Xu et al. 2015) |
| retinitis pigmentosa | p. R871C | -1.89 | p. W459* | (Xu et al. 2015) |
| retinitis pigmentosa | p. A974V | 0.30 | p. A418P | (Xu et al. 2015) |
| retinitis pigmentosa | p. L1399P | -8.41 | p. N633Sfs*10 | (Xu et al. 2015) |
| LCA | p. C329R | -1.05 | p. T484M | (Xu et al. 2015) |
| LCA | p. E790K | 0.00 | p. E522Gfs*6 | (Xu et al. 2015) |
| Atrioventricular septal defect | p. V108M | -2.59 | p. E1065K | (Priest et al. 2016) |
| LCA, renal failure | p. L440P | 0.04 | unknown | (Hull et al. 2016) |
| Retinal dystrophy with skeletal defects | p. E664K | -12.34 | p. E664K | (Khan, Bolz, and Bergmann 2014; Miller et al. 2013) |
| JATD | p. G212R | 1.59 | c.2399+1G>T | (Perrault et al. 2012) |
| JATD | p. V292M | -0.80 | p. G522E | (Schmidts et al. 2013) |
| JATD | p. V292M | -0.80 | p. N460Kfs28 | (Schmidts et al. 2013) |
| Mainzer-Saldino | p. G140R | -2.03 | p. E164* | (Schmidts et al. 2013) |
| Mainzer-Saldino | p. G212R | 1.59 | p. Ala1306Glyfs*56 | (Perrault et al. 2012) |
| Mainzer-Saldino | p. I233M | -1.35 | c.2399+1G>T | (Perrault et al. 2012) |
| Mainzer-Saldino | p. I233M | -1.35 | p. I233M | (Perrault et al. 2012) |
| Mainzer-Saldino | p. Y311C | 2.79 | p. Ile286Lysfs*6 | (Perrault et al. 2012) |
| Mainzer-Saldino | p. E664K | -12.34 | c.2399+1G>T | (Perrault et al. 2012) |
| Mainzer-Saldino | p. C1360R | 1.19 | c.2399+1G>T | (Schmidts et al. 2013) |
